# Prophylactic Treatment of Undernourished Mice with Cotrimoxazole Induces a Different Profile of Dysbiosis with Functional Metabolic Alterations

**DOI:** 10.1101/2021.11.01.466808

**Authors:** Lívia Budziarek Eslabão, Gabriela Farias Gubert, Lucas C. Beltrame, Isis M. A. de Mello, Oscar Bruna-Romero, Carlos R. Zárate-Bladés

**Author notes:** These authors contributed equally to this work (Lívia Budziarek Eslabão, Gabriela Farias Gubert). (L.B.E.); (O.B.R.); (G.F.G.); (L.C.B.); (I.M.A.d.M.); (C.R.Z.B.).

## Abstract

Childhood malnutrition affects physiology and development, it increases infection rate which may not present clinical signs in severe cases. To overcome this issue, the World Health Organization recommends prophylactic treatment with Cotrimoxazole (SXT) along with nutritional recovery. This treatment is controversial since evidence of reduction in morbidity and mortality is not a consensus and could induce the development of antibiotic resistant bacteria. Moreover, the impact of the use of this wide-spectrum antibiotic on gut microbiota, in a critical period of development and weakness, is unknown. To understand how SXT prophylaxis could affect gut microbiota in undernutrition, we induced protein-energy undernutrition in weaning C57BL/6 mice for three weeks and treated animals with SXT for two weeks. Using 16S rRNA gene sequencing we compared the taxonomic composition and metabolic pathways of control mice, animals submitted to undernutrition (UND), treated with SXT, or undernourished and SXT treated (UND+SXT). Undernutrition protocol was responsible for increasing Bacteroidetes and decreasing Firmicutes abundance. We identified that UND mice had a significant increase in predicted pathways related to metabolic syndromes later in life. The prophylactic SXT treatment alone resulted in significant loss in community richness and beta diversity. In addition, we identified the reduction of 6 families in SXT treated mice, including the butyrate producers Lachnospiraceae and Ruminococcaceae. The double challenge (UND+SXT) resulted in a reduction in Clostridiaceae family and in the urea cycle pathway, both related to the fermentation of amino acids, the intestinal epithelial permeability, and the healthy gut environment. Our results show that SXT prophylaxis during an undernourishment period in young mice did not re-establish the undernourished microbiota community composition similar to healthy controls, but induces a distinct dysbiotic profile, with functional metabolic consequences.

## 2. Introduction

Undernutrition still affects approximately 200 million children every year, been considered one of the leading underlying causes of morbidity and mortality according to the World Health Organization (WHO) [1]. Undernutrition is defined as the imbalance between the nutrients and/or energy ingestion and the individual’s basic needs to sustain the body’s homeostasis and its specific functions, and, in case of infants, the adequate growth [1]. In addition, the acute form of malnutrition during childhood affects several types of organs and functions, from bones to neuronal development, metabolism, immunity and even the gut microbiota [2–4].

In recent years, the microbiota has emerged as one of the major contributors to maintain the individual’s healthy status, performing a range of functions from nutrients’ me-tabolism, development and modulation of the immune system, direct protection against infections and even influences behavior and cognition [5]. The early gut microbiota can be greatly modulated by several factors during neonatal life, including mode of delivery, breastfeeding, use of antibiotics, environmental exposure, and nutritional status. Under-standing how nutrition and the gut microbiota of an individual interact with each other is extremely necessary to better comprehend the pathogenesis of undernutrition and to de-velop better prevention measures and more effective treatments [6].

Children suffering from malnutrition are more prone to infection, but they may not show signs of clinical infection [7]. As a result, WHO recommends as prophylactic treatment a course of Cotrimoxazole, a broad-spectrum antibiotic, in severe malnourished children [8]. The treatment is controversial, being classified based in weak evidence by WHO itself [9]. In a multicenter, double-blind randomized controlled trial, researchers found no increase in survival [10]. There is also the issue of global concern in increasing microbial resistance [11], as well as the potentiality of disruption of gut microbiota, resulting in broad-spectrum antibiotics-induced dysbiosis [12]. Nonetheless, there is also the opinion that the use of cotrimoxazole during undernutrition treatment could preserve the structure of the microbiome or even be beneficial for it since would avoid the invasion of pathogens during a period of particular vulnerability [13].

In this study we reproduced human infant undernutrition and its treatment in mice. The aim was to define the effects of undernutrition, cotrimoxazole or their association on microbiome composition and functionality. Our results show that each situation results in a different type of dysbiosis and that WHO’s malnutrition preconized treatment might not lead to the reestablishment of a healthy gut microbiome.

## 3. Materials and Methods

### Animal and study design

C57BL/6 mice aged 3-4 weeks of age were randomly housed in groups of 3 animals per microisolator cage (Alesco, Campinas, Brazil) on sterilized wood chip bedding under controlled temperature (21 ± 1 °C) and humidity (50 ± 20%) with a 12 h light/dark cycle. All animals were acclimated for 4 days to recover from transport stress before the onset of the experimental protocol. During acclimation period, mice had ad libitum access to sterilized distilled water and irradiated diet.

After acclimation, mice weighing 10.9 ± 1.52 g (mean ± SD) were randomly grouped into ad libitum control group or undernourished group. To study the effect of antibiotic therapy on food intake and gut microbiota, the control group and the undernourished group were divided into two sub-groups: no additional treatment and cotrimoxazole-treated. Thus, the experiment was carried out in four groups: control group (CON) (n = 6), cotrimoxazole-treated group (SXT) (n = 6), undernourished group (UND) (n = 6) and undernourished and cotrimoxazole-treated group (UND+SXT) (n = 6). Undernourishment model protocol was adapted from Mittal and Woodward [14]. Undernourished groups (UND and UND+SXT) were feed with diets containing only 60% of the total food consumed by CON and SXT groups, which equals to a total of 1.4 g of food per animal consumed daily by undernourished groups. Feeding was performed in the afternoon to re-spect the mice’s circadian rhythms. The food restriction protocol was sustained for three weeks.

The SXT protocol was carried out for two weeks prior the end of the food restriction period. Mice of SXT and UND+SXT groups were treated with SXT (a combination of 25 mg/kg of sulfamethoxazole + 5 mg/kg of trimethoprim), per animal daily for two weeks as recommended by WHO [8] for cases of severe malnutrition without apparent infection. SXT was purchased in a veterinary pharmacy with banana flavoring, for greater palatability to the animals. The drug application volume was adjusted to 50 μL, containing 10 μL cotrimoxazole and 40 μL of ultrapure water (Merck KGaA, Darmstadt, Germany). Finally, the drug was administered orally through a cannula and syringe system.

All procedures were performed in accordance with the Health Guide for the Care and Use of Laboratory Animals from Brazilian College of Animal Experimentation and were approved by the Institution’s Ethics Committee under protocol number 5560250219. All efforts were made to minimize animal suffering and to reduce the number of animals used in the experiments.

### Fecal sampling and DNA extraction

Fecal samples were collected from mice individually at day 0 (before the food restriction protocol), and at day 21 (end of the food restriction protocol). Samples were col-lected from each mouse independently by performing a tail-lift and aseptically collecting the fecal content direct from the anus into sterile tubes. Fecal samples were immediately transferred to liquid nitrogen and subsequently storage at −80 °C until processing.

DNA extraction was performed on weighted fecal samples using a FastDNA™ SPIN Kit (MP Biomedicals, Santa Ana, USA) according to the manufacturer’s recommendation.

### 16S rRNA gene sequence and analysis

The DNA extracts were quantified with Qubit dsDNA BR Assay Kit (Invitrogen™, Thermo Fisher Scientific, Carlsbad, USA) and amplified using primers 341F (5’-CCTAYGGGRBGCASCAG-3’) and 806R (5’-GGACTACNNGGGTATCTAAT-3’) for the V3-V4 hypervariable region of the 16S rRNA gene associated to a barcode sequence as de-scribed by Yu et al. [15]. Polymerase chain reactions were performed with 15 μL Phusion® High-Fidelity PCR Master Mix (New England Biolabs, Ipswich, USA), 0.2 μM of forward primer, 0.2 μM of reverse primer, and 10 ng of template DNA. Amplification conditions consisted in an initial denaturation at 98 °C for 1 min, followed by 30 cycles of denaturation at 98 °C for 10 s, annealing at 50 °C for 10 s, and extension 72 °C for 60 s, with a final extension 72 °C for 5 min. Amplification was confirmed through electrophoresis in agarose gel, resulting in amplicons with approximately 400-450 bp. For DNA library, amplicons were purified with Qiagen Gel Extraction Kit (Qiagen, Hilden, Germany) and prepared using TruSeq® DNA PCR-Free Sample Preparation Kit (Illumina, San Diego, USA) following manufacturer’s instruction. Finally, the library was sequenced on an Illumina HiSeq 2500 (Illumina, San Diego, USA), resulting in 250 bp paired-end reads. 16S rRNA gene sequence was performed by GenOne Biotech (Rio de Janeiro, Brazil).

Paired-end 16S rRNA gene sequence with low quality were filtered with Trimmomatic v0.36 [16] according to sequence size and Phred score. Nucleotides with a Phred score under 33 at the beginning and end of each sequence and sequences shorter than 200 nucleotides were considered low quality and removed. Barcode and primer sequence were also removed. Paired-end sequences were merged using DADA2 [17], available with the Quantitative Insights Into Microbial Ecology 2 (QIIME 2) software [18]. Chimera removal, singletons filtering, amplicon sequence variant generation (ASV), and rare ASV removal were also assessed using DADA2 pipeline. Operational taxonomic units (OTUs) were as-signed using the Greengenes v13.8 database [19] based on a 97% sequence similarity ac-cording to VSEARCH [20]. PICRUSt2 was used to predict metagenomic functions based on the normalized OTU tables [21].

### Statistical analysis of 16S rRNA sequencing data

All statistical analyses for 16S rRNA sequence data were performed in R v4.1.0 [22]. Alpha diversity measure of bacterial richness (observed species and Chao1) and diversity (Shannon and Simpson) were analyzed using phyloseq R package [23]. To test the statis-tical significance of alpha diversity, Student’s t-test or ANOVA followed by Tukey post-hoc test was applied for parametric data, and Mann-Whitney U test or Kruskal-Wallis followed by Dunn’s test of multiple comparisons for non-parametric data. Bray-Curtis distance metrics were used to access beta diversity through phyloseq R package [23] and vegan R package [24]. To access beta diversity statistical significance, a permutation multivariate analysis of variance (adonis) was conduct with 10000 permutations [25]. Principal Coordinates Analysis (PCoA) of Bray-Curtis distance was performed using vegan R package [24]. Differences in bacterial taxa abundance between experimental groups was evaluated using Kruskal-Wallis test and LEfSe (linear discriminant analysis (LDA) coupled with effect size measurements) analysis [26]. The LEfSe analysis was performed under the following conditions: the α value for the factorial Kruskal–Wallis test and pairwise Wilcoxon test among classes was < 0.05, and the threshold on the logarithmic LDA score for discriminative features was > 2.0. Predict metagenomic functions were analyzed through LEfSe and Kruskal-Wallis test with Benjamini–Hochberg False Discovery Rate (FDR) adjusted p-value. Data with p < 0.05 were considered to be significant.

## 4. Results

### 4.1. Changes in body weight and daily weight gain in C57BL/6 mice during undernourishment induction

C57BL/6 mice aged 3-4 weeks were randomly assigned to CON or UND groups (Day – 3). CON group mice had food access ad libitum. The undernourishment protocol consisted in a reduction of 40% of the daily food consumption for three weeks (from experimental day 0 to day 21) [14] (Figure 1A).

**Figure 1.**
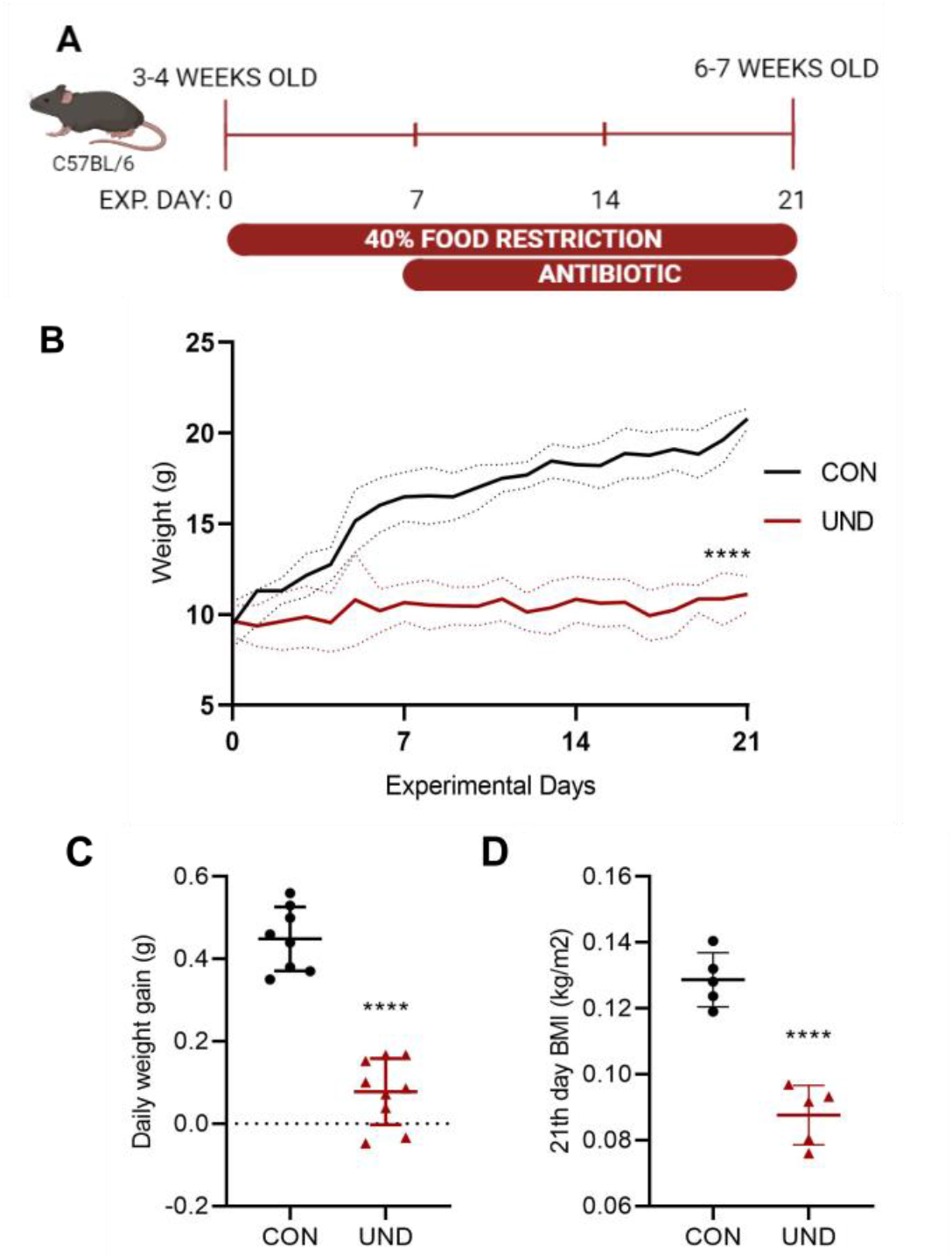
Changes in body weight during undernourishment induction through food restriction. (a) Experimental design, C57BL/6 mice with 3-4 weeks old were submitted to 40% food restriction for 21 experimental days (exp. day 21). (b) Body weight (g) changes in male C57BL/6 mice during the experimental protocol. (c) Daily weight gain (g) for the 3 weeks food restriction period. (d) BMI at the 21the experimental day. n = 5 animals per group. Experiments were repeated at least three times. Data for one typical experiment are shown. Statistical significances were assessed by Mann-Whitney U test. Data expressed as mean ± SD. **** p < 0.0001.

The undernourishment protocol resulted in a significant body weight loss when compared to same age healthy C57BL/6 mice (Figure 1B). The lowest body weight was observed at the end of the food restriction protocol with undernourished mice weighting 10.99 g (± 1.18) and control mice 19.62 g (± 1.28) (p < 0.0001). As expected, the daily weight gain in undernourished mice (0.088 g ± 0.080) was significantly lower when compared to the daily weight gain in control mice (0.449 g ± 0.078) (p < 0.0001) during the establishment of the food restriction protocol (Figure 1C). And at the experimental day 21, the end of food restriction, undernourished mice had a significantly lower body mass index (BMI) (0.088 kg/m2 ± 0.009) in relation to control (0.129 kg/m2 ± 0.008) (p < 0.0001)

### 4.2. Effects of undernourishment on the microbiota composition and function

To study the effects of undernourishment on the microbiota composition and func-tion, the gut microbiota was analyzed through 16S rRNA gene sequencing of fecal samples. After data processing and quality check, we observed a total of 2,879,810 reads with an average of 115,192 reads per sample. Sequences were clustered into OTUs with 97% identity, resulting in a total of 11,126 OTUs with an average of 445 OTUs per sample. We analyzed the fecal microbiota composition between CON and UND mice at the end of the food restriction period. Regarding the microbiota composition, 17 main phyla were found in which Bacteroidetes, Firmicutes, and Proteobacteria were the most abundant (Figure 2A). Alpha diversity analysis showed there was no difference in community richness (Observed OTU and Chao1) and diversity indices (Shannon and Simpson) between CON and UND mice (Data not showed). To illustrate intra-group microbial community’s differences after the undernourishment protocol, PCoA was performed using Bray-Curtis distances, revealing distinct findings (Figure 2B). UND microbiota was significantly different in community composition when compared to CON microbiota (adonis with 10000 permutations, p = 0.004)

**Figure 2.**
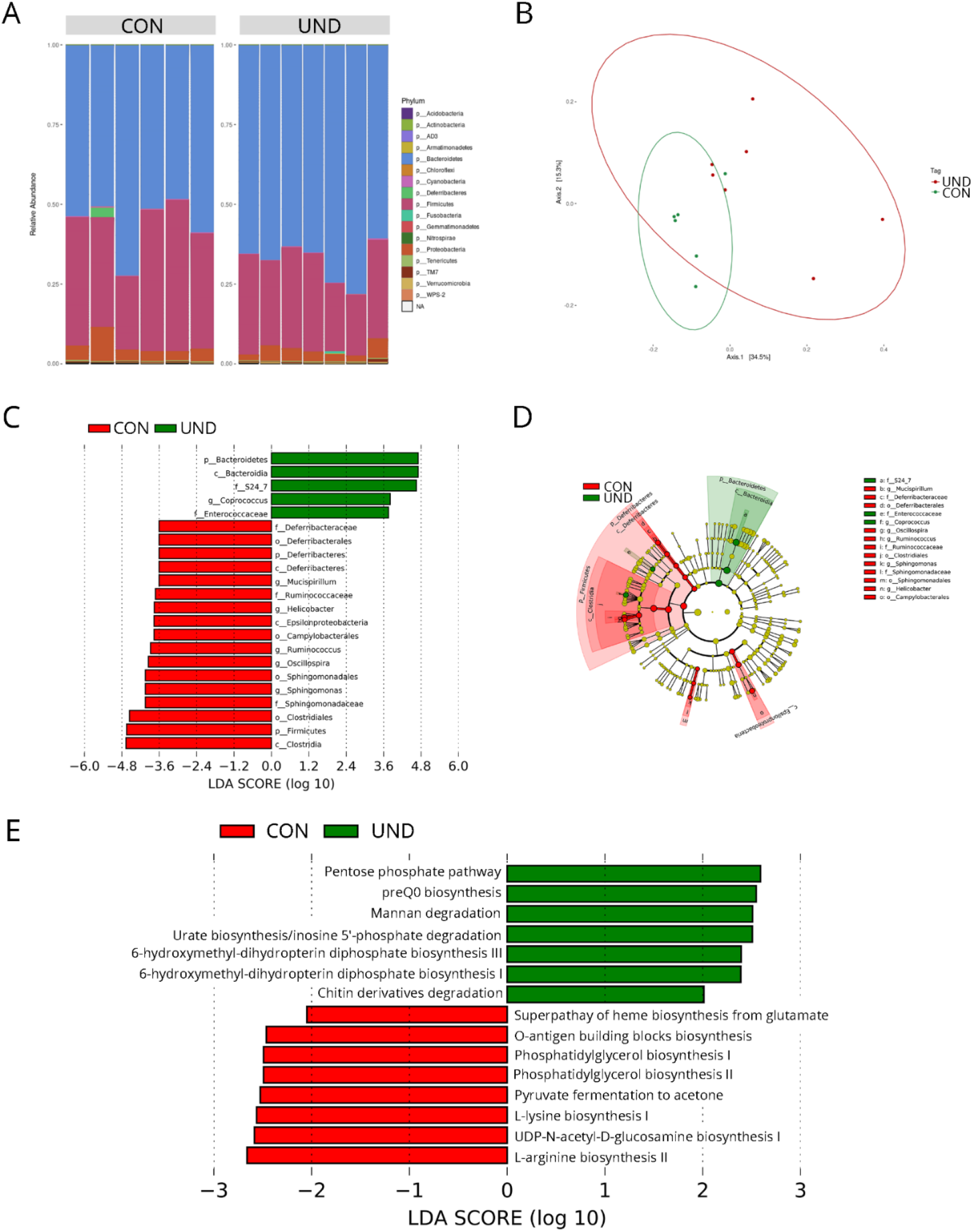
The gut microbiota composition and inferred functional content of gut microbiota between control (CON) and undernourished (UND) C57BL/6 mice. (a) Relative abundance at the phylum level for groups of CON (n = 6) and UND (n = 7) C57BL/6 mice; (b) Principal coordinate analysis (PCoA) of Bray Curtis distances among groups of CON (n = 6) and UND (n = 7) C57BL/6 mice. Each point corresponds to a community from a single mouse. Colors indicate group identity. Ellipses show the 95% confidence intervals. Intra-group differences were evaluated using adonis test (p < 0.05); (c) LefSe linear discriminant analysis (LDA) scores showing differentially abundant taxonomic clades with an LDA score > 2.0 in the gut microbiota of CON (n = 6) and UND (n = 7) C57BL/6 mice; (d) Cladogram of LefSe linear discriminant analysis (LDA) scores for significantly different taxonomic clades for the gut microbiota of CON (n = 6) and UND (n = 7) C57BL/6 mice; (e) LDA scores showing significant pathway differences between CON (n = 6) and UND (n = 7) C57BL/6 mice. Significant differences between groups were tested with Kruskal-Wallis test (p < 0.05).

LefSe analysis was used to compare the gut microbiota taxa that were significantly different between CON and UND groups (Figure 2C). When comparing taxa at phylum level, UND mice showed an increase in Bacteroidetes and a decrease in Firmicutes and Deferribacteres. In addition, UND mice also presented an increase in Coprococcus genus (0.0056 ± 0.002 vs. 0.002 ± 0.0003) and a decrease in Mucispirillum (0.0009 ± 0.0008 vs. 0.007 ± 0.012), Helicobacter (0.0062 ± 0.002 vs. 0.0168 ± 0.011), Ruminococcus, Oscillospira (0.0299 ± 0.005 vs. 0.0440 ± 0.011), and Sphingomonas (0.0001 ± 0.0001 vs. 0.0005 ± 0.0001) genera. Figure 2D shows differences in the abundance of taxonomic clades of LDA score > 2.0 between CON and UND mice.

To determine whether the observed taxonomic differences between groups played a role in function, metabolic pathways were accessed using PICRUSt2. LefSe analysis compared metabolic changes in the gut microbiota in each group (Figures 2D), all displayed pathways presented a LDA score > 2.0. The UND group changes were mainly related to cellular growth. Pathways related to the biosynthesis of sugar nucleotides [O-antigen building blocks biosynthesis (0.066 ± 0.004 vs. 0.073 ± 0.005), UDP-N-acetyl-D-glucosamine biosynthesis I (0.052 ± 0.006 vs. 0.062 ± 0.007)], amino acids [L-lysine biosynthesis I (0.074 ± 0.006 vs. 0.083 ± 0.006), L-arginine biosynthesis II (0.081 ± 0.009 vs. 0.092 ± 0.006)], cell membrane lipids [phosphatidylglycerol biosynthesis I (0.0921 ± 0.003 vs. 0.099 ± 0.006), phosphatidylglycerol biosynthesis II (0.0921 ± 0.003 vs. 0.099 ± 0.006)] were less present when compared to CON group. In addition, the compromising of growing related metabolic pathways, UND presented increases in pathways of degradation of cell components, such as carbohydrates [pentose phosphate pathway (0.073 ± 0.008 vs. 0.062 ± 0.005)], polysaccharides [mannan degradation (0.047 ± 0.007 vs. 0.038 ± 0.005)], purine nucleotides [urate biosynthesis/inosine 5’-phosphate degradation (0.112 ± 0.005 vs. 0.101 ± 0.008)], and cellular wall [chitin derivatives degradation (0.00011 ± 0.00022 vs. 0 ± 0)]. There was also an increase in vitamin biosynthesis-related pathways of preQ0 biosynthesis (0.067 ± 0.007 vs. 0.05 ± 0.006), 6-hydroxymethyl-dihydropterin diphosphate biosynthesis III (0.097 ± 0.005 vs. 0.088 ± 0.007), and 6-hydroxymethyl-dihydropterin diphosphate biosynthesis I (0.096 ± 0.005 vs. 0.087 ± 0.007).

### 4.3. Effects of Cotrimoxazole (SXT) on the gut microbiome

To study the effect of SXT treatment on the microbiota composition and function, we analyzed the fecal microbiota composition between CON and SXT mice groups at the end of the antibiotic treatment. Regarding the microbiota composition, the same 17 main phyla found in UND were present in SXT (Figure 3A). Alpha diversity analysis showed significant difference in community richness (Observed, p = 0.0397; and Chao1, p = 0.04061) between CON and SXT mice (Figure 3B). In contrast, there was no difference in diversity indices (Shannon and Simpson) between both groups (Figure 3B). To illustrate microbial community changes after the antibiotic treatment, PCoA was performed using Bray-Curtis distances (Figure 3C). Following antibiotic treatment, SXT microbiota is sig-nificantly different in community composition when compared to CON microbiota (ADONIS, p = 0.02).

**Figure 3.**
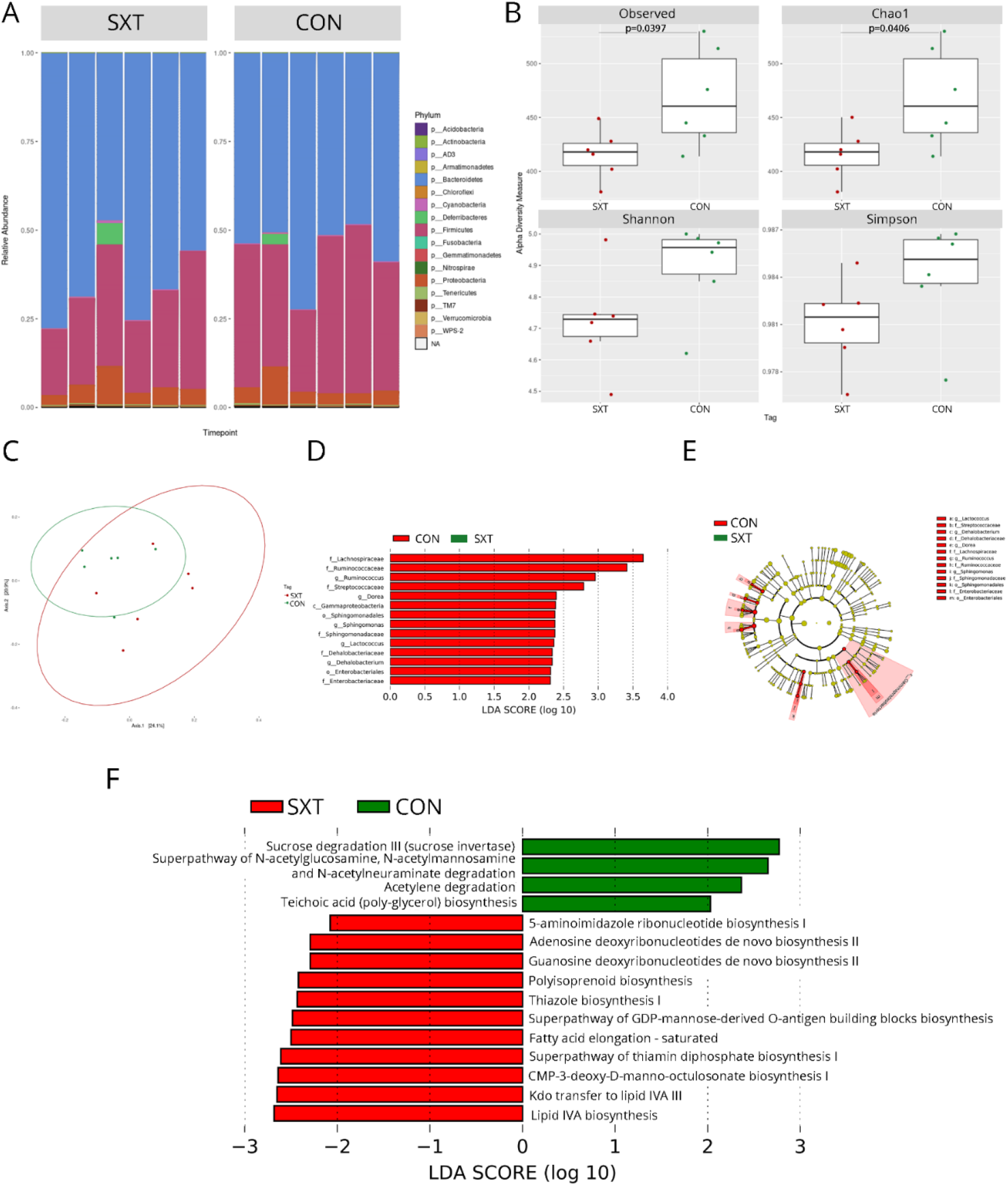
The gut microbiota composition and inferred functional content of gut microbiota between of control (CON) and cotrimoxazole (SXT) C57BL/6 mice. (a) Relative abundance at the phylum level for groups of CON (n = 6) and SXT (n = 6) C57BL/6 mice; (b) Richness (Observed and Chao1) and diversity (Shannon and Simpson) indexes for groups of CON (n = 6) and SXT (n = 6) C57BL/6 mice; (c) Principal coordinate analysis (PCoA) of Bray Curtis distances among groups of CON (n = 6) and SXT (n = 6) C57BL/6 mice. Each point corresponds to a community from a single mouse. Colors indicate group identity. Ellipses show the 95% confidence intervals. Intra-group differences were evaluated using adonis test (p < 0.05); (d) LefSe linear discriminant analysis (LDA) scores showing differentially abundant taxonomic clades with an LDA score > 2.0 in the gut microbiota of CON (n = 6) and SXT (n = 6) C57BL/6 mice; (e) Cladogram of LefSe linear discriminant analysis (LDA) scores for significantly different taxonomic clades for the gut microbiota of CON (n = 6) and SXT (n = 6) C57BL/6 mice; (f) LDA scores showing significant pathway differences between CON (n = 6) and SXT (n = 6) C57BL/6 mice of PICRUSt predicted relative MetaCyc pathways abundances. Significant differences between groups were tested with Kruskal-Wallis test (p < 0.05).

LefSe showed the presence of 14 taxa in CON microbiome when compared to SXT microbiome (Figure 3D). We did not identify significant taxa presence in SXT microbiome. Differences did not appear at a phylum level but, 6 families were impacted by antibiotic treatment, Lachnospiraceae, Ruminococcaceae, Streptococcaceae, Sphingomonadaceae, Dehalobacteriaceae, and Entereobacteriaceae (Figure 3D). Figure 3E shows differences in the abundance of taxonomic clades of LDA score > 2.0 between CON and SXT mice.

Figure 3E showed the heatmap plot and LefSe analysis comparing metabolic changes in the gut microbiota of CON and SXT mice. SXT increased pathways related to nucleotide synthesis [5-aminoimidazole ribonucleotide biosynthesis I (0.113 ± 0.003 vs. 0.1104 ± 0.003); adenosine deoxyribonucleotides de novo biosynthesis II (0.09398 ± 0.002 vs. 0.093 79 ± 0.002); guanosine deoxyribonucleotides de novo biosynthesis II (0.09398 ± 0.002 vs. 0.093 79 ± 0.002); superpathway of GDP-mannose-derived O-antigen building blocks biosynthesis (0.072 ± 0.003 vs. 0.068 ± 0.003); CMP-3-deoxy-D-manno-octulosonate biosynthesis I (0.0627 ± 0.003 vs. 0.0553 ± 0.005)], vitamin synthesis [thiazole biosynthesis I (0.0501 ± 0.003 vs. 0.0457 ± 0.004); superpathway of thiamin diphosphate biosynthesis I (0.0803 ± 0.003 vs. 0.0743 ± 0.004)], lipid metabolism [fatty acid elongation (0.098 ± 0.001 vs. 0.095 ± 0.002)], and alcohols [polyisoprenoid biosynthesis (0.083 ± 0.001 vs. 0.08 ± 0.001)]. In contrast, SXT decreased carbohydrate and amine degradation [sucrose degradation III (0.055 ± 0.006 vs. 0.0724 ± 0.012); superpathway of N-acetylglucosamine, N-acetylmannosamine and N-acetylneuraminate degradation (0.0403 ± 0.003 vs. 0.0536 ± 0.008)], fermentation of pyruvate or short-chain fatty acids [acetylene degradation (0.0244 ± 0.004 vs. 0.031 ± 0.004)], and formation of cell wall [teichoic acid biosynthesis (0.006 ± 0.001 vs. 0.009 ± 0.001)] when compared to CON mice.

### 4.4. Impact of the combination of undernourishment and the use of cotrimoxazole (UND+SXT) on microbiota composition and function

The analysis of the microbiota composition showed the same 17 main phyla found in CON were also pre-sent in UND-SXT (Figure 4A). The undernutrition treated with a prophylactic course of antibiotic did not alter alpha diversity (data not showed). However, UND+SXT microbiota is significantly different in community composition when compared to CON microbiota (adonis, p = 0.0003) (Figure 4B).

**Figure 4.**
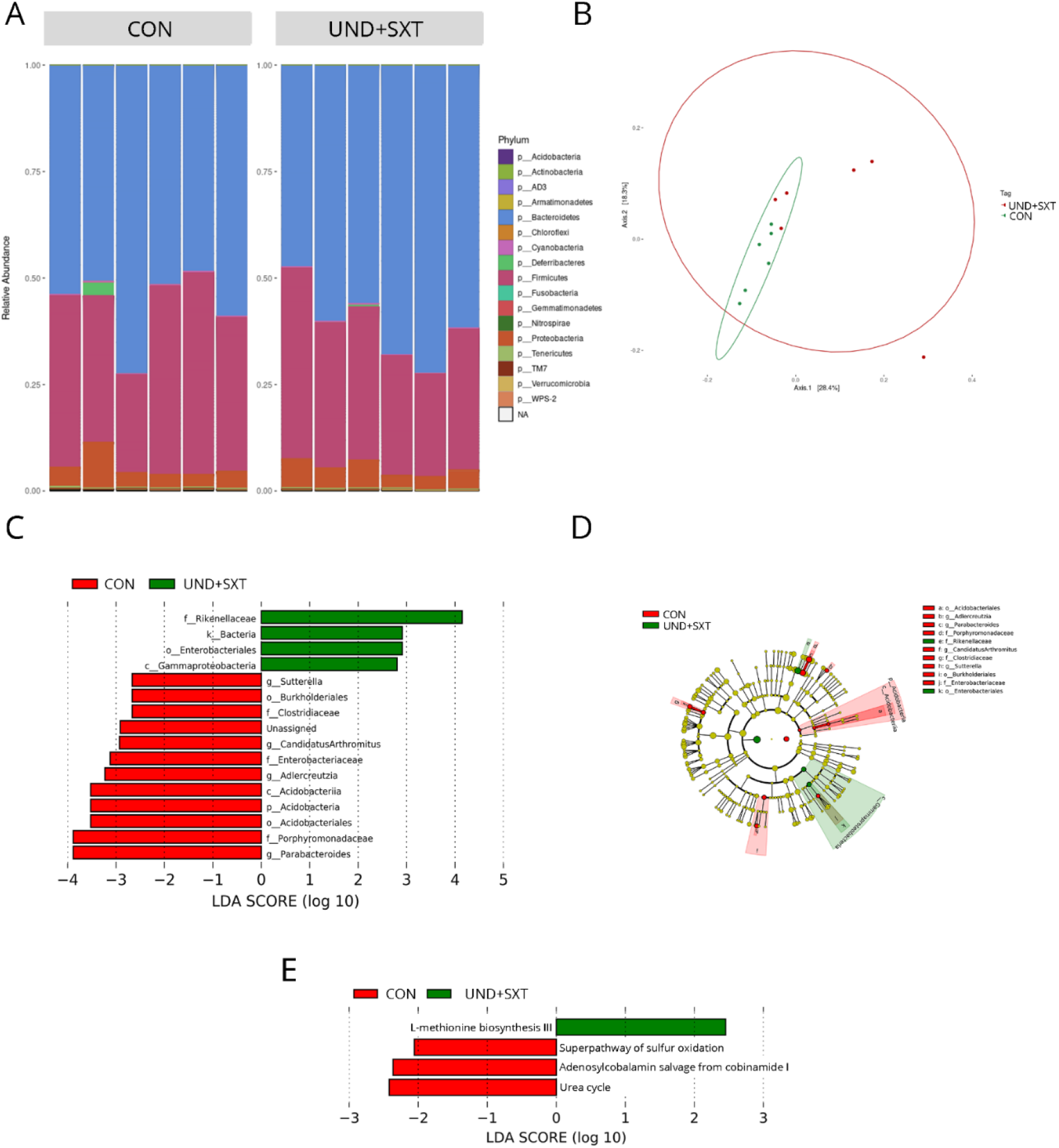
The gut microbiota composition and inferred functional content of gut microbiota between of control (CON) and cotrimoxazole-treated undernourished (UND-SXT) C57BL/6 mice. (a) Relative abundance at the phylum level for groups of CON (n = 6) and UND+SXT (n = 6) C57BL/6 mice; (b) Principal coordinate analysis (PCoA) of Bray Curtis distances among groups of CON (n = 6) and SXT (n = 6) C57BL/6 mice. Each point corresponds to a community from a single mouse. Colors indicate group identity. Ellipses show the 95% confidence intervals. Intra-group differences were evaluated using adonis test (p < 0.05); (c) LefSe linear discriminant analysis (LDA) scores showing differentially abundant taxonomic clades with an LDA score > 2.0 in the gut microbiota of CON (n = 6) and UND+SXT (n = 6) C57BL/6 mice; (d) Cladogram of LefSe linear discriminant analysis (LDA) scores for significantly different taxonomic clades for the gut microbiota of CON (n = 6) and UND+SXT (n = 6) C57BL/6 mice; (e) LDA scores showing significant pathway differences between CON (n = 6) and UND+SXT (n = 6) C57BL/6 mice of PICRUSt predicted relative MetaCyc pathways abundances. Significant differences between groups were tested with Kruskal-Wallis test (p < 0.05).

LefSe showed a significant reduction in UND+SXT microbiome when compared to CON (Figure 4C). Clostridiaceae, Enterobacteriaceae, and Porphyromonadaceae family were significantly impacted by being reduced in the double treatment, while Rikenellaceae were increased. Besides that, four genera are significantly lower in UND+SXT when compared to CON, Parabacteroidetes (0.0407 ± 0.009 vs. 0.0584 ± 0.0119), Candidatus arthromitus (0.00033 ± 0.00019 vs. 0.00119 ± 0.00065), Suterella (0.00028 ± 0.00009 vs. 0.00083 ± 0.0004), and Adlercreutzia (0.000116 ± 0.00007 vs. 0.00022 ± 0.00006). Figure 4D shows differences in the abundance of taxonomic clades of LDA score > 2.0 between CON and UND+SXT mice.

The LefSe analysis comparing metabolic changes in the gut microbiota exposed changes in the double treatment are presented in Figure 4E. The microbiota of UND+SXT mice presented a reduction in pathways related to enzymes cofactor synthesis [adenosylcobalamin salvage from cobinamide I (0.336 ± 0.037 vs. 0.364 ± 0.041)], sulfur and nitrogen compounds metabolism [superpathway of sulfur oxidation (0.052 ± 0.005 vs. 0.068 ± 0.014); urea cycle (0.217 ± 0.02 vs. 0.253 ± 0.018)] when compared to CON group. And a unique pathway increase, in amino acids synthesis [L-methionine biosynthesis III (0.392 ± 0.035 vs. 0.313 ± 0.034)].

### 5.5. The use of cotrimoxazole during undernourishment in infant mice does not revert undernutrition effects on microbiota, but results in a distinct profile of dysbiosis

The microbiota composition keeps the same 17 main phyla observed in all other treatment groups (Figure 5A). The undernutrition treated with a prophylactic course of cotrimoxazole did not alter alpha diversity when compared to UND and CON (data not showed). However, CON, UND, and UND+SXT microbiota profiles are significantly different in community composition when compared to each other (adonis, p = 0.00029) (Figure 5B).

**Figure 5.**
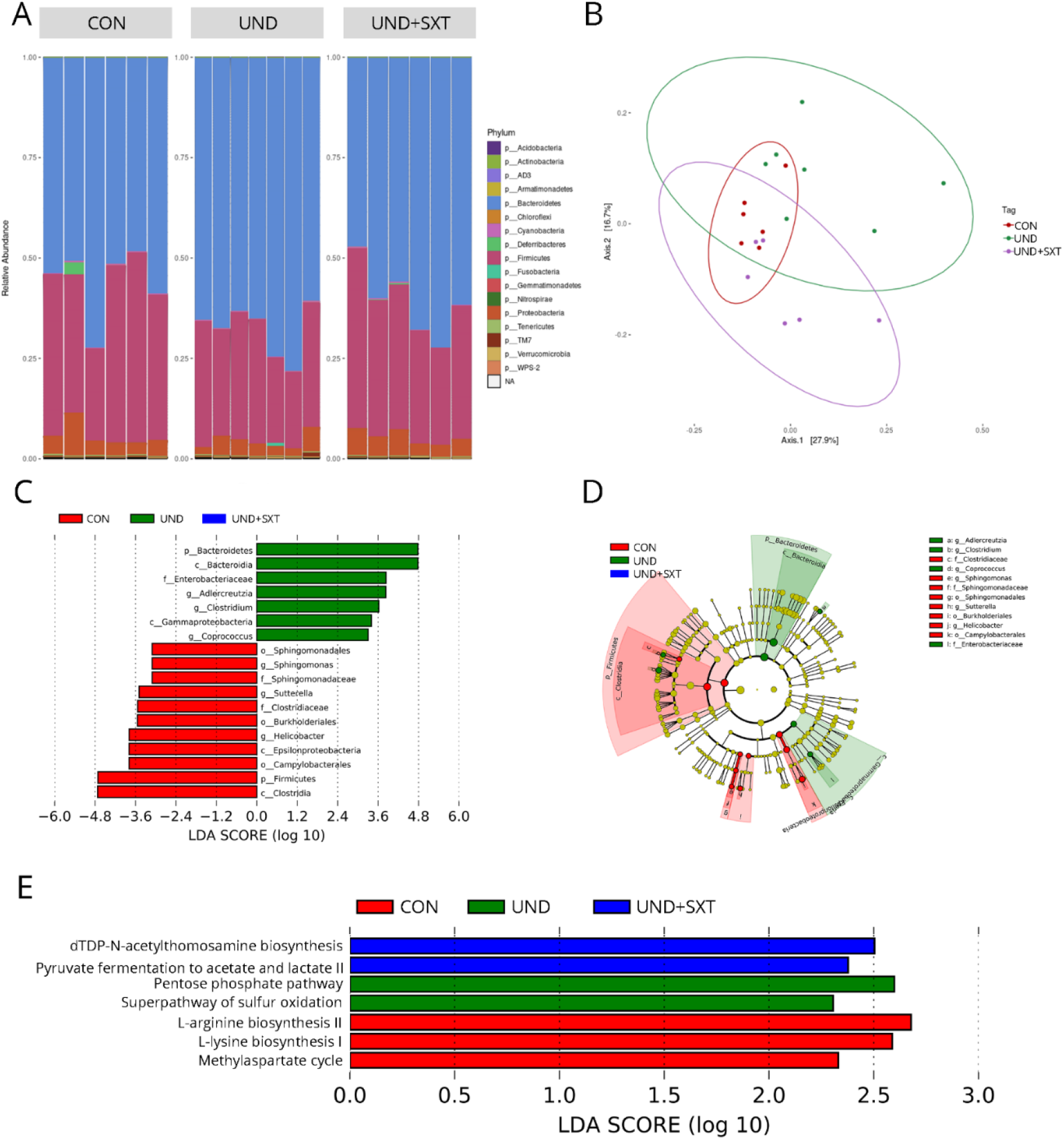
The gut microbiota composition and inferred functional content of gut microbiota between of control (CON), undernourished (UND), and cotrimoxazole-treated undernourished (UND-SXT) C57BL/6 mice. (a) Relative abundance at the phylum level for groups of CON (n = 6), UND (n = 7), and UND+SXT (n = 6) C57BL/6 mice; (b) Principal coordinate analysis (PCoA) of Bray Curtis distances among groups of CON (n = 6), UND (n = 7), and SXT (n = 6) C57BL/6 mice. Each point corresponds to a community from a single mouse. Colors indicate group identity. Ellipses show the 95% confidence intervals. Intra-group differences were evaluated using adonis test (p < 0.05); (c) LefSe linear discriminant analysis (LDA) scores showing differentially abundant taxonomic clades with an LDA score > 2.0 in the gut microbiota of CON (n = 6), UND (n = 7), and UND+SXT (n = 6) C57BL/6 mice; (d) Cladogram of LefSe linear discriminant analysis (LDA) scores for significantly different taxonomic clades for the gut microbiota of CON (n = 6), UND (n = 7), and UND+SXT (n = 6) C57BL/6 mice; (e) LDA scores showing significant pathway differences between CON (n = 6), UND (n = 7), and UND+SXT (n = 6) C57BL/6 mice of PICRUSt predicted relative MetaCyc pathways abundances. Significant differences between groups were tested with Kruskal-Wallis test (p < 0.05).

LefSe showed that both CON and UND had significant differences in the microbiome composition when the three groups are taken into account (Figure 5C). Although, UND+SXT microbiota did not show differences in the presence of specific taxa when compared to both other groups, the taxa identified as differentially present in the other two groups by LefSe indicate that these taxa are less abundant in UND+SXT mice. Although Bacteroidetes phylum was significantly increased in UND mice when compared to CON and UND+SXT mice, we observed a significant increase in members of Firmicutes (Clostridium and Coprococcus) and Actinobacteria (Adlercreutzia) phyla. Besides that, CON mice presented a significant increase in Firmicutes phylum. We also observed an increase in Helicobacter, Sutterella, and Sphingomonas genera, members of Proteobacteria phylum. Figure 5D shows differences in the abundance of taxonomic clades of LDA score > 2.0 between CON, UND and UND+SXT mice.

The LefSe analysis comparing metabolic changes in the gut microbiota of control mice, undernourished mice, and double treatment mice are presented in Figure 5E. When compared to CON and UND mice, the microbiota of UND+SXT mice presented a reduction in pathways related to energy production from organic substrates [methylaspartate cycle (0.00001 ± 0.00001 vs. 0.00006 ± 0.00006 vs. 0.00009 ± 0.00005 for UND+SXT, UND, and CON, respectively)] and inorganic nutrient metabolism [superpathway of sulfur oxidation (0.02 ± 0.002 vs. 0.034 ± 0.012 vs. 0.027 ± 0.0064 for UND+SXT, UND, and CON, respectively)]. UND+SXT microbiota also presented an increase in pathways related to sugar nucleotide biosynthesis [dTDP-N-acetylthomosamine biosynthesis (0.071 ± 0.008 vs. 0.052 ± 0.011 vs. 0.065 ± 0.017 for UND+SXT, UND, and CON, respectively)] and short-chain fatty acids fermentation [pyruvate fermentation to acetate and lactate II (0.332 ± 0.005 vs. 0.329 ± 0.012 vs. 0.325 ± 0.009 for UND+SXT, UND, and CON, respectively)]. In addition, UND microbiota presented an increase in the generation of precursor metabolites and energy pathway [pentose phosphate pathway (0.155 ± 0.024 vs. 0.187 ± 0.027 vs. 0.152 ± 0.018 for UND+SXT, UND, and CON, respectively)]. Furthermore, CON microbiota presented an increase in pathways related to amino acid biosynthesis [L-lysine biosynthesis I (0.196 ± 0.009 vs. 0.189 ± 0.011 vs. 0.202 ± 0.012 for UND+SXT, UND, and CON, respectively) and L-arginine biosynthesis II (0.225 ± 0.014 vs. 0.207 ± 0.017 vs. 0.227 ± 0.013 for UND+SXT, UND, and CON, respectively)].

## 5. Discussion

In the present study, we characterized the gut microbiota profile of undernourished (UND), cotrimoxazole-treated (SXT), and undernourished cotrimoxazole-treated (UND+SXT) C57BL/6 mice after 3 weeks of protein-energy undernourishment and compare them to healthy control mice (CON). Our experimental protocol was carried out in animals just after weaning, intended to resemble undernutrition in children. Moreover, the main objective of our study was to stablish the effects of SXT on the microbiota of undernourished mice, a wide-spectrum antibiotic therapy recommended by WHO and routinely used in undernourished children.

First of all, our results showed that C57BL/6 mice who underwent 3 weeks of protein-energy undernourishment did not present significant differences in fecal microbiota richness compared to healthy controls but, they did present a significant change in beta diversity analysis. These are in accordance with the results of [27]. We observed major phylum-level alterations in UND mice compared to CON, including a depletion in obligate anaerobic bacteria, such as Firmicutes (Ruminococcaceae and Oscillospiraceae) and Deferribacteres (Deferribacteraceae). These findings support clinical studies where the in-fant malnutrition was associated with a reduction on anaerobic bacteria [28–30]. Other studies have associated the presence of Oscillospiraceae with leanness and a lower body mass index (BMI) in children, adults, and germ-free mice colonized with human fecal samples [31,32]. However, Oscillospiraceae decreases in abundance during the onset of inflammatory diseases, such as Inflammatory bowel disease (IBD) [33], and the under-nourished state in early life has been associated with increased inflammation markers [34,35]. The microbiota immaturity led by a poor nutrition might result in the increase of enteropathogenic species and the dysfunction of the gut barrier, factors associated with inflammation on early life [36]. Thus, considering these studies alongside with our findings related to the decrease in Oscillospiraceae in UND mice, allow to speculate that our undernourished model might also compromise the gut barrier, leading to an inflammatory state and the further alteration in the gut microbiota. In addition, we observed that Bacteroidetes, mostly from the Muribaculaceae family, were significantly increased in UND mice compared to healthy controls. These findings contradict other studies in which a decrease in Bacteroidetes were observed in undernourished children and undernour-ished neonatal mice [30,37], although in the last case, the authors used outbred CD1 mice and not C57BL/6 isogenic mice as here. Previous studies demonstrated that Muribaculaceae was correlated with the inner mucus layer formation and function in the colon, and with the production of propionate, a short-chain fatty acid (SCFA) [38,39]. Thus, further histological analysis is needed to understand the effects of malnutrition on the intestinal tissue of this model.

We also observed differences in metabolic pathways between CON and UND mice. Pathways involved in the biosynthesis and/or the degradation of carbohydrates, polysaccharides, purine nucleotides, and cellular wall were increased in UND mice. The double burden of malnutrition (DBM) is characterized by the coexistence of undernutrition in early life and overweight, obesity, and noncommunicable diseases related to diet in later life [40]. Several studies have associated undernutrition at young age to disease later in life, such as diabetes, hypertension, and metabolic syndrome [40,41]. In addition, metabolic syndromes are also correlated with hyperuricemia, a pathological condition characterized by the overproduction and/or decreased excretion of uric acid [42]. Although not yet fully elucidated, evidence indicates that gut bacteria directly affect host urate degradation [43]. In our study, we identify an increase in the purine nucleotides degradation pathway, which leads to the urate biosynthesis, suggesting a possible on-set of hyperuricemia, leading to metabolic dysfunctions later in life. We also observed that UND mice had an increase in the pentose phosphate pathway, a metabolic pathway involved in the glucose oxidation. The pentose phosphate pathway is increased in the gut microbiome after the depletion of body glycogen stores during periods of insufficient carbohydrate consumption [44]. We may speculate that the pentose phosphate pathway was favored in our UND mice in response to carbohydrate consumption limitation during the food restriction protocol. Furthermore, our study showed that UND mice might increase the metabolic pathways leading to the folate (vitamin B9) biosynthesis. Folate is essential for the maintenance of Treg cells [45], and deficiencies in vitamin B9 lead to the development of intestinal inflammation [46]. It may be the case that the higher abundance of folate pathways in UND mice microbiota when compared to CON mice is a mechanism to regulate the inflammatory environment in consequence of malnutrition. Finally, UND mice lost representation of heme biosynthesis, sugar nucleotide biosynthesis, phosphatidylglycerol biosynthesis, pyruvate fermentation, L-lysine biosynthesis, and L-arginine biosynthesis pathways. Lysine levels are decreased in metabolic syndrome patients and it correlates negatively with cardio-metabolic features and inflammatory biomarkers [47]. Our findings further suggest that malnutrition during infancy in mice might lead to metabolic alterations later in life.

Despite the efforts to achieve better clinical outcomes, 10 to 15% of undernourished children are not able to recover even after controlled treatment [48]. Several studies have reported a higher prevalence of clinically significant infections among children who have been hospitalized for severe malnutrition [49,50]. However, the diagnose of severe infection is difficult during undernutrition, since undernourished children might not present clinical signs of infection [49]. Nonetheless, the WHO recommends the use of SXT as treatment for severe undernourished infants together with the ready-to-use therapeutic food (RUTF), even in cases with no detected infection. To evaluate the impact of SXT in the gut microbiota of undernourished infant mice, we first characterized the antibiotic-driven disruption of this wide-spectrum antibiotic in healthy infant mice. We observed that SXT mice had a significant loss in community richness (Observed and Chao1), but no alterations in diversity indices (Shannon and Simpson). SXT prophylaxis has been widely studied in HIV exposed infants [51,52], and in this case, SXT therapy did not altered the microbiome taxonomic composition or functional metabolic pathways [51]. Other studies with HIV and hematological patients did not find neither a significant difference in α-diversity in the gut microbiome after SXT prophylaxis [52,53]. Nonetheless, SXT treatment resulted in a microbiota community profile distinct from healthy control animals according to beta diversity analysis. We confirmed these dysbiotic profile through LefSe analysis, in which we identify 6 bacterial families that were depleted by SXT: Lachnospiraceae, Ruminococcaceae, Streptococcaceae, Sphingomonadaceae, Dehalobacteriaceae, and Enterobacteriaceae. We also observed a reduction in *Dehalobacterium*, *Dorea*, and *Sphingomonas* genera in SXT mice. Reduction in Lachnospiraceae family is a highly profound short-term effect on human gut microbiome after administration of commonly used antibiotics, such as β-lactams and fluoroquinolones [54]. Lachnospiraceae and Ruminococcaceae family members are important butyrate producers [55]. Butyrate is a main gut mucosal immune regulator derived from the microbiota and one of the best functional markers for a healthy mature gut microbiota [55]. In models for autism spectrum disorder, studies showed that Dehalobacterium decrease is associated with altered behavior, an increase of TNF-α expression, the onset of a colon proinflammatory state in female mice and increased gut permeability in male mice [56]. As mentioned, we also observed a reduction in Sphingomonas, a major environmental microorganism that is not found in high abundance in the gut microbiota [57]. However, its presence has been shown to be able to stimulate iNKT cells [57,58]. Similarly, we also detected a decrease of Enterobacteriaceae family in SXT treated mice. Several pathogenic species responsible for major economic loss and health-related impacts are part of this family [59]. In addition, high abundance Enterobacteriaceae’s family members were correlated with neutrophilia and lower oral vaccine responses in a Bangladeshi infant cohort with a 4-8% prevalence of moderate wasting malnutrition and a 10-12% prevalence of moderate stunting [60]. The use of antibiotic therapy usually results in the depletion of butyrate-producing bacteria, which leads to the reduction in the expression of the receptor mediating butyrate oxidation [61,62]. This alteration is followed by lower epithelial oxygen utilization and increased gut oxygen availability which promotes the expansion of aerobic bacteria including Enterobacteriaceae members [62]. Even though we detected a reduction in taxa related to butyrate production, we did not detect an increase in Enterobacteriaceae colonization after SXT prophylaxis. In a remarkable study, Kau et al. [6] demonstrated a link among weight loss associated phenotype, disruption of the gut barrier function and the development of an abnormal mucosal immunity to the presence of Enterobacteriaceae family members and related enteropathogenic microorganisms, in gnotobiotic mice after colonization with fecal microbiota from Malawian twins discordant for kwashiorkor. Taken together, our results suggest that SXT prophylaxis favors a gut microenvironment with fewer taxa responsible for immune system stimulation in addition to impair the colonization of opportunistic pathogens.

We also observed the purine nucleotide biosynthesis pathway increased in SXT mice. Commensal bacteria are responsible for the constitutive development of intestinal Th17 cells differentiation, due in part through purinergic receptor signaling [63]. On the other hand, immunosupressive and anti-inflammatory effects can also be mediated by purinergic signaling. The purine nucleotide biosynthesis pathway 5-aminoimidazole ribonucleotide biosynthesis and its variants, were identified in lower abundance in IBD and colorectal cancer, indicating that the downregulation of this purinergic signaling contribute to gut inflammation [63,64].

The superpathway of GDP-mannose-derived O-antigen building blocks biosynthesis was elevated in SXT mice microbiome. O-antigen is part of the bacteria outer membrane lipopolysaccharide [65], indicating elevated LPS levels in the gut. Another molecule that constitutes the LPS is Lipid A [65]. The superpathway of (Kdo)2-lipid A biosynthesis was elevated in SXT mice, supporting the idea of Gram-negative bacteria increase in STX treated mice. LPS produced by healthy gut microbiota has important role for immunotolerance of the microbial community and it was reduced in SXT mice [66]. In addition, we also observed a reduction in the superpathway of N-acetylglucosamine, N-acetylmannosamine and N-acetylneuraminate degradation. When deprived of dietary fiber, the gut microbiota uses the mucus glycoproteins as a nutrient source, resulting in the degradation of the mucus layer and in the access of gut epithelial to lumen pathogens [67]. Thus, the SXT therapy might mitigate the colonic mucus barrier erosion in undernourished individuals through the reduction of the superpathway of N-acetylglucosamine, N-acetylmannosamine and N-acetylneuraminate degradation.

Long-term cotrimoxazole prophylaxis is well recognized with clinical benefits in HIV infection, including the reduction of morbidity and mortality of HIV-positive children and adults in areas with serious infectious threats [52,68]. Bourke et al. [52] showed that cotrimoxazole reduces systemic inflammation in HIV-positive children through microbiota alteration, together with reducing immune and epithelial cell activation. Although gut microbiota modifications related to cotrimoxazole therapy are available in the literature, SXT effects in the undernourished gut microbiota and in the undernourished gut metabolic functions are still missing. To elucidate the impact of SXT in the gut of undernourished mice, we compared undernourished mice treated with a prophylactic course of this antibiotic to healthy control mice. We identified a significant decrease in the abundance of Sutterella and Adlercreutzia in UND+SXT mice when compared to CON group. Sutterella has been associated with pro-inflammatory cytokines in digestive disorders [69]. In contrast, the presence of Adlercreutzia is usually associated with the restoration of health benefits in the microbiota due to its role in SCFA production and antioxidants metabolism [70]. SXT prophylaxis also demonstrated alterations related to the predicted metabolic pathways in undernourished mice. Compared with CON, fecal microbiota of UND+SXT presented reduction in superpathway of sulfur oxidation. Altering this metabolic pathway has both, beneficial and harmful effects for the host [71,72]. Accumulating hydrogen sulfide (H2S) has been linked with colonic inflammation and conditions such as colorectal cancer [73], inflammatory bowel disease (IBD) and ulcerative colitis (UC) [71]. At the same time, mice who had inhibition of H2S synthesis presented mucosal injury and inflammation in the small intestine and colon [72], suggesting the importance of the presence of H2S at appropriate levels. The possible H2S accumulation could be the result of the reduction of sulfur oxidation superpathway, which could explain the increase in the L-methionine biosynthesis III pathway, observed in the same animals [74].

The UND+SXT mice also showed a reduction in the urea cycle, which intermediate metabolites could also impair intestinal epithelial barrier function [75]. The urea cycle is responsible for the conversion of metabolic wastes from amino acid catabolism in ammonia, and then into urea [76]. Clostridiaceae family (including Clostridium) is the major responsible for both the fermentation of amino acids into urea as well as the hydrolysis of urea into ammonia. The products of amino acids fermentation will act later as a further support for Clostridiaceae enrichment in a health gut [77]. The ammonia is then used in amino acid biosynthesis while its excess is eliminated in the feces. All these pathways’ synergy help mammals to remove urea and promote homeostasis [76]. Our results showed a reduced presence of the Clostridiaceae family in UND+SXT animals, indicating the alteration of a healthy gut environment in these animals.

LefSe analysis of CON vs UND vs UND-SXT mice groups, showed a similar trend of Firmicutes previously observed in humans, that is undernourished children showed a lower percentage of Firmicutes in their gut microbiota when compared to health children [30]. Antibiotic treatment has also been correlated with the reduction of Firmicutes abundance [54] as we also observed in our experiments.

The reduction of amino acids biosynthesis pathway in both UND and UND+SXT animals when compared to healthy controls probably reflects the lack of food intake, which impacts cell growth. But this lack of aminoacids synthesis could also impact host physiology as up to 20% of circulating plasma lysine, body protein lysine, and urinary lysine derived from microbial sources [78]. Besides being a precursor for protein biosynthesis [79], lysine regulates other amino acids synthesis pathways, such as arginine [80] which was also downregulated. Both, lysine and arginine have been found reduced in stunted children [81]. At the same time, UND microbiota increases pentose phosphate pathway, which could suggest a strategy to deal with oxidative stress using NADPH generation for detoxification [82]. When undernourished animals received SXT, the dTDP-N-acetylthomosamine biosynthesis increased, which is the pathway responsible for producing a glycolipid common to all members of the Enterobacteriaceae [83]. Malnourished children in nutritional recovery have been reported to increase the number of antibiotic-resistant Enterobacteriaceae after amoxicillin therapy [84]. The same phenomenon could possibly be happening in our model.

Our study is the first to evaluate the effects of SXT prophylaxis in the undernourished gut microbiota of young animals. Nonetheless, it was not exempt of important limitations. These include the evaluation of local and systemic inflammatory markers and their correlation to several observations made on the microbiome modulation of our experimental groups. In addition, a longer time point of evaluation would bring information about the duration of the dysbiosis observed in each protocol.

## 6. Conclusions

Altogether our results present the impact of the prophylactic cotrimoxazole treatment on gut microbiota and possible consequences in host physiology in a murine model of childhood undernutrition. The controversial treatment alters gut microbiota in a different way than the undernutrition or SXT alone, creating a third dysbiotic profile that alters metabolic pathways related to amino acids synthesis and energy production. Other studies will be necessary to understand if this affects enterocytes metabolism, intestine permeability, or how these taxonomic changes relate to the host immune system.

## Author Contributions

L.B.E. and G.F.G., designed the study, handled the animals, processed samples, analyzed the data, and wrote the manuscript. L.C.B. handled the animals and processed samples. I.M.A.d.M. handled the animals. O.B.R. and C.R.Z.B. designed the study, supervised the work, write the article and revised the manuscript. All authors read and approved the final version of the manuscript.

## Funding

No direct funding for the execution of this study was received. L.B.E., G.F.G., and I.M.A.d.M received a CAPES (Coordenação de Aperfeiçoamento de Pessoal de Nível Superior) student fellowship.

## Institutional Review Board Statement

The study was conducted according to the guidelines of the Health Guide for the Care and Use of Laboratory Animals from Brazilian College of Animal Experimentation, and approved by the Institutional Ethics Committee of Universidade Federal de Santa Catarina, UFSC (protocol code 5560250219, approved on July 9^th^, 2019).

## Informed Consent Statement

Not applicable.

## Data Availability Statement

The data presented in this study are available on request from the corresponding author.

## Acknowledgments

We are grateful to all members of the Laboratory of Applied Immunology and of the Laboratory of Immunoregulation of UFSC.

## Conflicts of Interest

The authors declare no conflict of interest.

## Notes

### Competing Interest Statement

The authors have declared no competing interest.

